# TRIM37 recognizes a bipartite degron to ubiquitinate centrosome substrates

**DOI:** 10.64898/2025.12.19.695442

**Authors:** Weronika E. Stachera, Judith Tafur, Nicole E. Familiari, Kan Yaguchi, Jeffrey B. Woodruff

## Abstract

Dysregulation of the E3 ubiquitin ligase TRIM37 is associated with tumor formation and Mulibrey nanism, a recessive developmental syndrome. TRIM37 regulates steady-state levels of centrosome proteins and limits their ectopic assembly, but how it recognizes and ubiquitinates its substrates is poorly understood. We found that TRIM37 directly ubiquitinates the centrosome-forming protein Cep192 at 7 lysines clustered near its C-terminus. TRIM37 binds Cep192 at a C-terminal intrinsically disordered region followed by an ASH domain (IDR+ASH8). Mutation of the 7 lysines or the IDR+ASH8 domain increased Cep192 levels and stability in cells, indicating loss of TRIM37-based regulation. Fusing IDR+ASH8 to an unrelated protein (GFP-EB1) was sufficient to enable its degradation via TRIM37. Biochemical assays revealed that IDR+ASH8 is primarily monomeric and binds TRIM37 via two separate coiled-coil motifs with mid-nanomolar affinity. We propose that the IDR+ASH8 motif is a bipartite degron for TRIM37, enabling it to target centrosome proteins and adjust their levels.

## INTRODUCTION

Centrosomes are major microtubule-organizing centers (MTOCs) in mitotic cells and are essential for accurate chromosome segregation. They are composed of two barrel-shaped centrioles surrounded by a supramolecular mass of protein called pericentriolar material (PCM) (Woodruff et al., 2014). To safeguard genome stability, cells must tightly control not only centriole duplication but also the abundance and organization of the PCM (Watanabe et al., 2019). Even modest disruptions in PCM protein levels can impair spindle assembly, promote aneuploidy, and drive tumorigenesis (Levine et al., 2017; Lingle et al., 2002; Xie et al., 2022). Although PCM composition and architecture are well understood, mechanisms that control the levels of core PCM scaffolding proteins remain poorly defined.

One mechanism for controlling PCM protein abundance is proteasome-mediated degradation. TRIM37, an E3 ubiquitin ligase and a key regulator of centrosome composition, has been shown to influence stability of several PCM components including Cep192, a conserved scaffolding protein required for PCM assembly (Balestra et al., 2021; Meitinger et al., 2020; Yeow et al., 2020). TRIM37 overexpression reduces cellular Cep192 levels, whereas loss of TRIM37 leads to the accumulation of ectopic cytoplasmic Cep192-containing foci (Meitinger et al., 2020). Whether TRIM37 targets Cep192 directly or indirectly through other pathways has not been tested. Furthermore, the molecular mechanism underlying Cep192 recognition and ubiquitination by an E3 ligase remains unknown.

Understanding this pathway is particularly important given the strong disease relevance of TRIM37. The genomic locus harboring *TRIM37* is amplified in several cancers— including neuroblastoma, breast, liver, and kidney tumors (Bhatnagar et al., 2014; Jiang et al., 2015; Meitinger et al., 2020; Miao et al., 2021)—and elevated TRIM37 levels contribute to chemoresistance (Chen et al., 2020; Nishibeppu et al., 2021; Przanowski et al., 2020; Tan et al., 2021; Yeow et al., 2020). Several studies proposed that excessive TRIM37 activity destabilizes PCM proteins, thereby compromising mitotic fidelity, which could contribute to cancer progression (Balestra et al., 2021; Meitinger et al., 2020; Yeow et al., 2020; Zhou et al., 2024). Conversely, loss-of-function mutations in *TRIM37* are the only known cause of Mulibrey nanism (MUL), an autosomal recessive growth disorder (Avela et al., 2000). Fixed fibroblasts derived from MUL patients displayed ectopic centrosome-like foci (Balestra et al., 2021; Meitinger et al., 2021) that contain Centrobin, a suspected substrate of TRIM37. Ectopic centrosome-like assemblies in MUL cells can act as aberrant MTOCs and induce chromosome mis-segregation, defects thought to contribute both to impaired growth and the elevated cancer risk associated with the disease.

TRIM37 contains a disordered C-terminal tail preceded by three conserved modules at its N-terminus (Wang et al., 2017): (i) a RING domain required for E2 recruitment and ubiquitin transfer, (ii) a B-box domain that mediates higher order assembly, and (iii) a MATH domain, recently implicated in dimerization and recognizing pathological Centrobin-rich foci (Bellaart et al., 2025; Yeow et al., 2025) (Fig. S1A). While TRIM37-mediated ubiquitination of Centrobin aggregates has been linked to multimerization-dependent activation, this mechanism appears specialized for targeting pathogenic assemblies rather than endogenous PCM substrates. Thus, the principles by which TRIM37 identifies and regulates physiological PCM scaffold proteins remain unresolved.

Here, we identified Cep192 as a direct target of TRIM37 and show that TRIM37 binds a composite region containing an intrinsically disordered region and a structured beta barrel domain (IDR+ASH8). We mapped 7 lysine residues that TRIM37 ubiquitinates to promote Cep192 degradation and demonstrate that the IDR+ASH8 module functions as a transferable degron to enable TRIM37-dependent degradation of unrelated proteins. Together, these findings reveal the molecular basis of Cep192 regulation and illustrate how a RING E3 ligase recognizes both unstructured and folded substrate elements to achieve substrate specificity.

## RESULTS

### TRIM37 directly ubiquitinates 7 lysines on the C-terminus of Cep192

While previous studies have shown that TRIM37 associates with Cep192 and regulates its cellular levels (Meitinger et al., 2020; Tkach et al., 2022; Yeow et al., 2020), there is no clear evidence that TRIM37 directly targets Cep192 for ubiquitination. Thus, we established an *in vitro* ubiquitination assay. We purified full-length Cep192 tagged with GFP (MW = 307 kDa) from baculovirus-infected insect cells (Fig. S1B) and combined it with E1 (UBE1) and E2 (UBCH5B) enzymes, ATP, MgCl_2_, and HA-tagged ubiquitin. We then incubated the reactions at 37°C for 1 hr, either in the presence or absence of purified TRIM37 (Fig. 1A). Western blotting revealed increased ubiquitination signal on high molecular weight proteins in the reaction containing TRIM37 (Fig. 1A).

**Figure 1.**
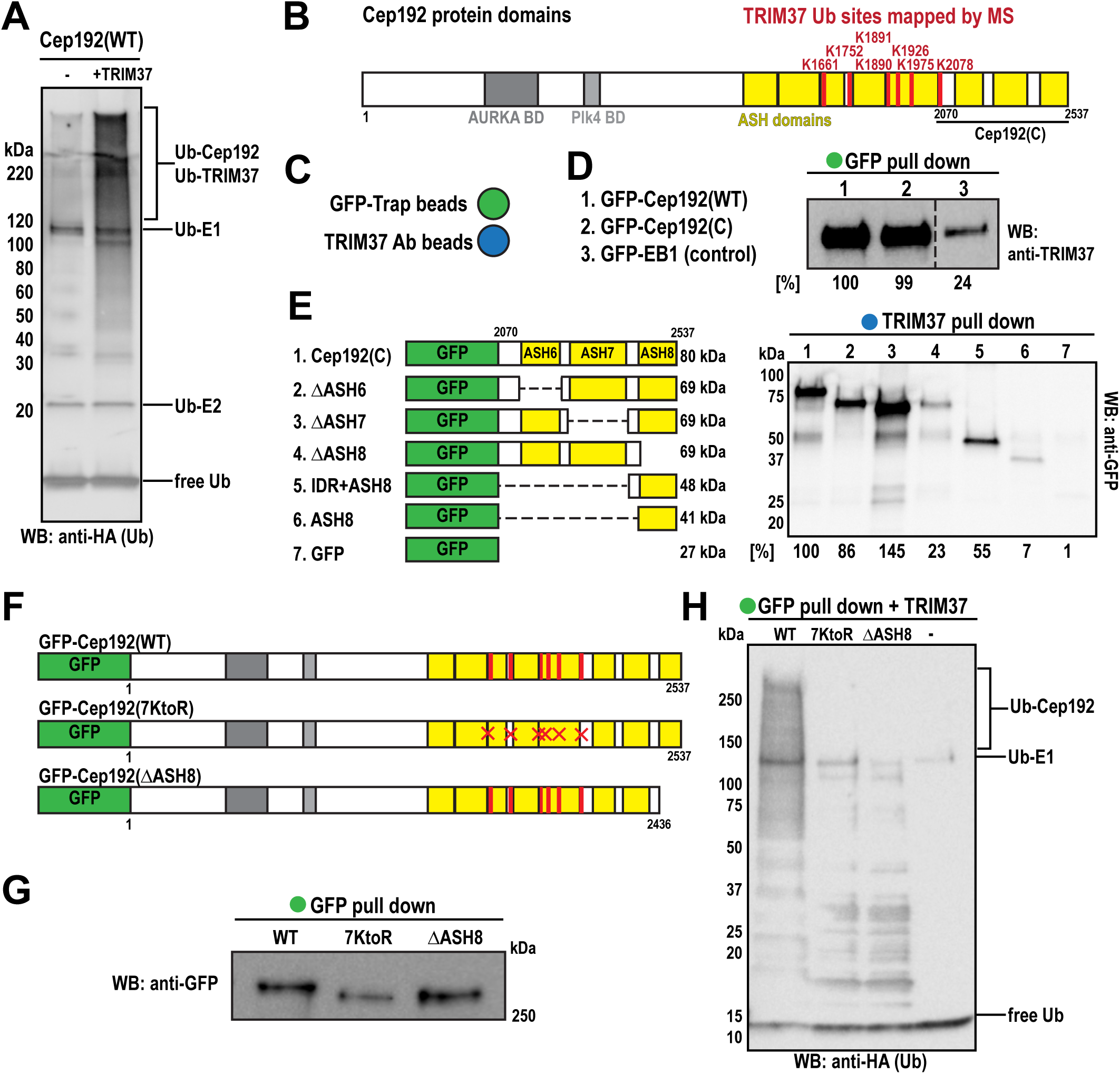
TRIM37 directly binds the IDR+ASH8 domain of Cep192 and ubiquitinates it on seven specific residues. A) Western blot against HA-tagged ubiquitin from reconstituted ubiquitination reactions performed without or with TRIM37. B) Schematic of Cep192 domains indicating ubiquitinated residues identified by mass spectrometry. C) Graphic representation of beads used for immunoprecipitation assays. Green: GFP-Trap magnetic particles; blue: beads conjugated to TRIM37 antibody. D) Western blot showing TRIM37 binding to Cep192 constructs. Percentages below represent relative binding capacity normalized to full-length Cep192. E) Western blot against GFP showing Cep192 fragments that bind TRIM37. Percentages below represent binding capacity normalized to the Cep192 C-terminal fragment. F) Cep192 constructs expressed in DLD-1 cells. G) Western blot against GFP after pull-down of Cep192 constructs from DLD-1 cells. H) Western blot against HA-tagged ubiquitin from reconstituted ubiquitination reactions using the Cep192 constructs purified in panel G.

To identify specific ubiquitination sites on Cep192, we used mass spectrometry to detect di-glycine (GG) remnants indicative of lysine ubiquitination. We identified seven modified sites that were significantly more abundant in the presence of TRIM37: K1661, K1752, K1890, K1891, K1926, K1975, and K2078 (Fig. 1B, S1C). Notably, these ubiquitination sites cluster near the C-terminal region of Cep192 previously shown to pull-down TRIM37 (Fig. 1B) (Meitinger et al., 2020). AlphaFold3 predicts that this region contains three beta barrel ASH-like domains connected by intrinsically disordered regions (IDRs) (Fig. 1B). We compared the efficiency of TRIM37 binding between full-length Cep192 and the C-terminus fragment (2070-2537 aa), both GFP-tagged (Fig. 1B). EB1-GFP served as a negative control. We incubated each GFP fusion protein with an equal amount of TRIM37 at 37°C for 1 hr, then precipitated them using GFP-Trap beads (Fig. 1C). Minimal TRIM37 was detected in the control, while both full-length and C-terminal Cep192 showed similarly strong binding (Fig. 1C, D). These results demonstrate that TRIM37 directly binds the Cep192 C-terminus and ubiquitinates it on 7 lysines.

### TRIM37 recognizes a bipartite motif at the C-terminus of Cep192

To define which elements mediate TRIM37 binding, we generated GFP-tagged Cep192 C-terminal truncations lacking individual ASH-like domains (ΔASH6, ΔASH7, and ΔASH8) (Fig. 1E, S1B). In pull-down assays with bead-immobilized TRIM37 (Fig. 1C), only Cep192(ΔASH8), which lacks the final 100 amino acids, failed to bind TRIM37 (Fig. 1E). Surprisingly, ASH8 alone did not bind strongly to TRIM37, indicating that additional elements are required for the interaction (Fig. 1E). Inclusion of a short IDR adjacent to the ASH8 domain restored TRIM37 binding (Fig. 1E). Collectively, these data identify the ASH8 domain, plus its preceding IDR (IDR+ASH8) as the primary recognition site for TRIM37.

To determine whether the 7 identified ubiquitination sites and the TRIM37-binding domain are required for Cep192 regulation, we generated human colorectal cells expressing GFP-tagged wild-type Cep192 (WT), a mutant in which all 7 lysines were replaced with arginines (7KtoR), or a mutant lacking ASH8 (ΔASH8) (Fig. 1F, S1D). We then lysed the cells, purified the proteins using GFP-Trap beads (Fig. 1C, G), and subjected them to the *in vitro* ubiquitination assay described in Fig. 1A. As expected, only Cep192(WT) was ubiquitinated by TRIM37, whereas no ubiquitination was detected for the 7KtoR or ΔASH8 mutants (Fig. 1H). We also tested if these Cep192 mutants bind TRIM37. As expected, the mutant lacking the ASH8 domain failed to pull down TRIM37, whereas WT and 7KtoR could, although 7KtoR binding was reduced compared to WT (Fig. S1E). Thus, mutating either the ubiquitin acceptor sites or the docking module prevents Cep192 ubiquitination by TRIM37.

### TRIM37-mediated ubiquitination controls Cep192 levels in human cells

We next asked whether this modification is necessary for Cep192 degradation in cells. We induced expression of each transgene with tetracycline, then washed out the tetracycline and measured the decay of the GFP-Cep192 signal over time. (Fig. 2A). Imaging showed that each construct localized to fluorescent foci that predominantly represent centrosomes (Fig. 2B, C), as verified by co-localization with a centriole marker (Fig. S2A). Notably, foci formed by both mutants were on average brighter than those formed by Cep192(WT) (Fig. 2D). RT-qPCR confirmed that mRNA expression was similar for all constructs (Fig. S1D), indicating that any changes in protein abundance are likely due to differences in degradation rates. Two days after tetracycline removal, GFP-Cep192(WT) foci were mostly undetectable, indicating efficient degradation (Fig. 2B–D). In contrast, foci made by both Cep192 mutants persisted with minimal intensity reduction (Fig. 2B–D). Western blotting further confirmed that the mutant proteins remained stable after tetracycline washout (Fig. 2E). Together, these findings demonstrate that TRIM37-driven Cep192 ubiquitination is essential for Cep192 turnover in cells.

**Figure 2.**
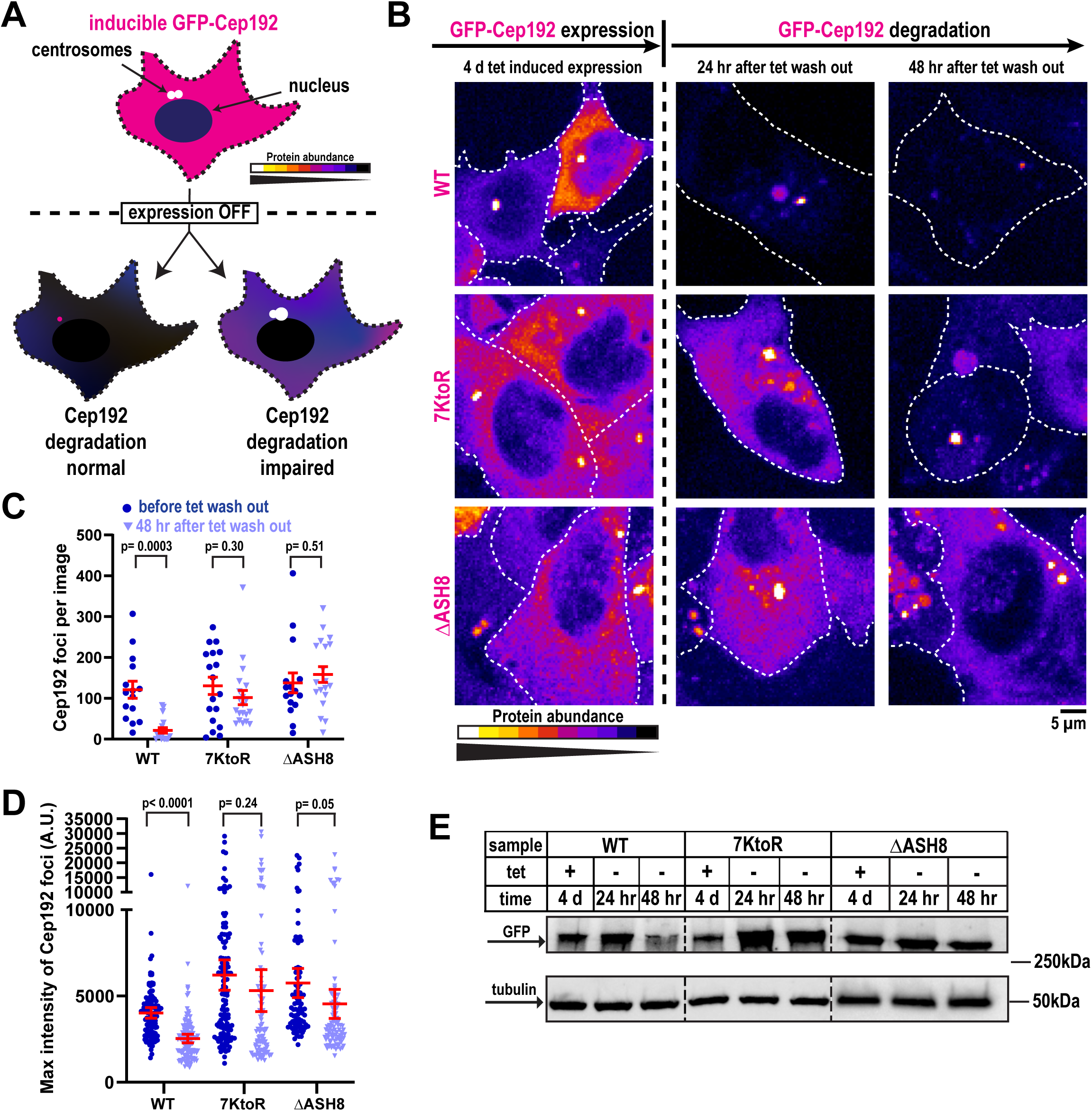
Ubiquitination-deficient Cep192 mutants show elevated steady-state levels and stability. A) Schematic of the construct degradation assay. GFP-Cep192 constructs localize to centrosomes and the cytoplasm after tetracycline-induced expression. After tetracycline washout, construct degradation is monitored; lighter color indicates higher protein abundance. B) Representative images showing degradation efficiency of each construct. C) Quantification of Cep192 foci per image (mean ± SEM; n ≥ 15 images per condition, 3 biological replicates) during expression (dark purple) and degradation (light purple) phases. P values from unpaired T-tests with Welch’s correction. D) Quantification of maximum fluorescence intensity of Cep192 foci (mean ± 95% CI; n > 90, 3 biological replicates) during expression (dark purple) and degradation (light purple) phases. P values from unpaired T-tests with Welch’s correction. E) Western blot against GFP showing total construct abundance under each condition. Tubulin detection was used as a loading control.

### Expression of degradation-resistant Cep192 causes mitotic arrest and reduced proliferation in cells

We next examined the functional consequences of impaired TRIM37-mediated degradation of Cep192. To assess cell viability, we expressed GFP-tagged WT, 7KtoR, or

ΔASH8 Cep192 in cells and monitored their growth over six days (Fig. 3A). Cells expressing the 7KtoR mutant showed a ∼50% reduction in viability compared to WT, whereas ΔASH8 expression resulted in a ∼75% decrease. To investigate potential causes of this phenotype, we performed live-cell imaging of synchronized cells released from late G2 arrest and tracked mitotic division (Fig. 3B). On average, mitotic division lasts ∼40 min in DLD-1 cells, but the duration can vary considerably, extending in some cases up to 2 hr (Bloomfield et al., 2020). For this reason, we imaged cells for up to 3 hr and classified cells that completed division within 110 min of mitotic entry as undergoing timely mitotic progression. ∼85% of cells expressing Cep192(WT) completed mitosis within 110 min, compared with ∼25% of 7KtoR-expressing and ∼38% of ΔASH8-expressing cells (Fig. 3C). Thus, cells expressing mutants showed a significantly longer mitotic duration, which may explain their reduced viability and proliferation. All cells still formed bipolar spindles (Fig. S2B), suggesting that the mitotic arrest was likely not caused by spindle assembly defects. We also noticed that Cep192 mutants exhibited reduced fluorescence intensity at mitotic centrosomes in comparison to Cep192(WT) (Fig. 3D). This stands in contrast to cells in interphase (Fig. 2). Thus, although ubiquitin-deficient Cep192 mutants are more abundant and accumulate more strongly at interphase centrosomes compared with wild-type Cep192, their accumulation at mitotic centrosomes is impaired.

**Figure 3.**
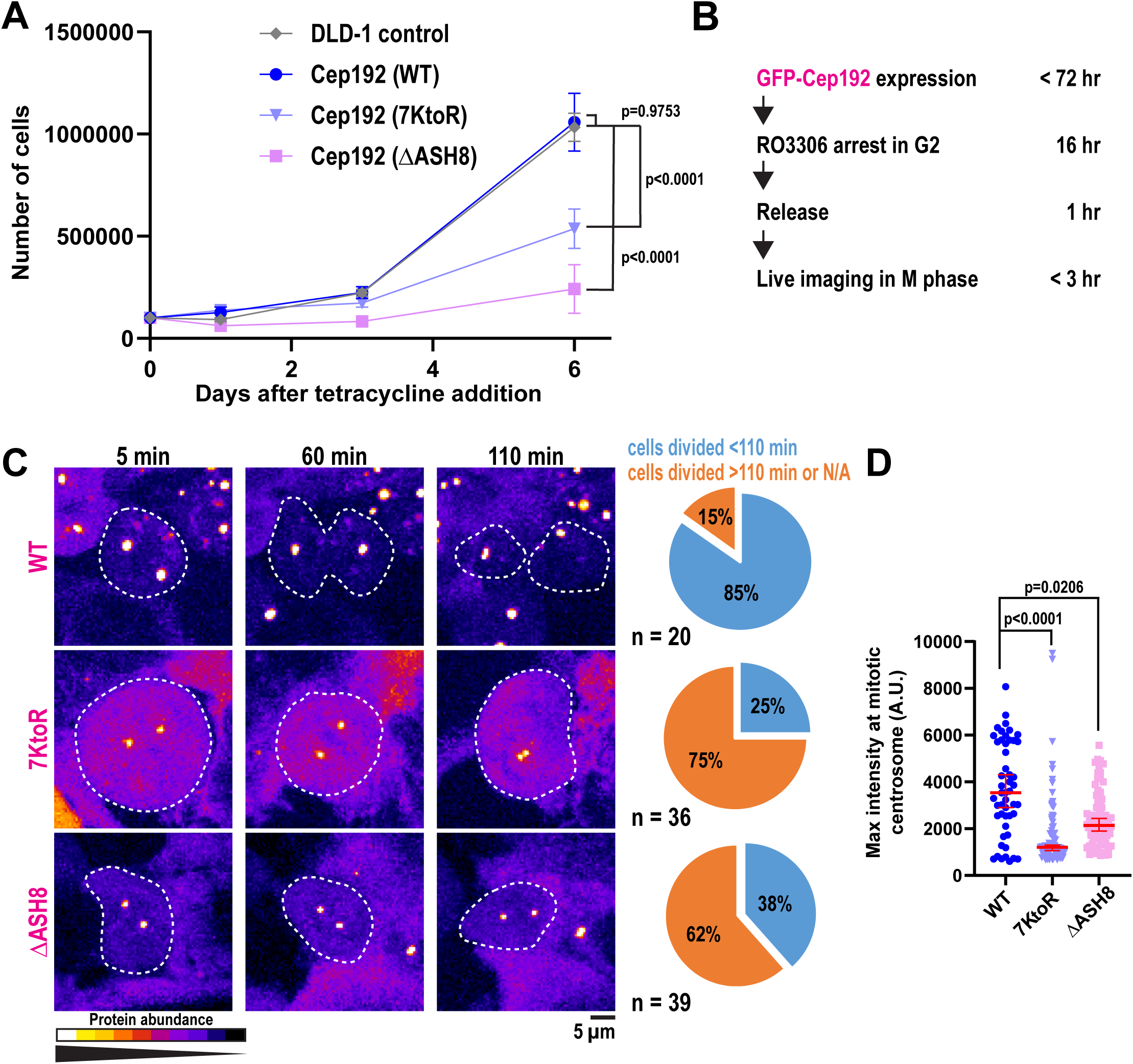
Cells expressing ubiquitination-deficient Cep192 mutants arrest in mitosis. A) Quantification of cell proliferation during expression of Cep192 constructs (mean ± 95% CI; n = 6 biological replicates). P values (day 6 viability) from one-way ANOVA followed by Dunnett’s multiple comparison test. B) Schematic showing the cell-synchronization timeline used before mitotic tracking. C) Representative images of cells in mitosis over time. Cells are outlined in white. Pie charts indicate the percentage of cells completing division within 110 min during imaging. D) Quantification of maximum fluorescence intensity of GFP-Cep192 constructs at mitotic centrosomes (median ± 95% CI; ≥ 50 centrosomes, 4 biological replicates). P values from Kruskal-Wallis test, followed by Dunn’s multiple comparison test.

### Adding the IDR+ASH8 domain to EB1 transforms it into a TRIM37 substrate

We next examined whether the IDR+ASH8 domain is sufficient for TRIM37 targeting in cells. The microtubule end-binding protein EB1 localizes to both microtubules and centrosomes (Berrueta et al., 1998; Bu & Su, 2001; Louie et al., 2004); however, it does not interact with TRIM37 in either proximity-based assays (Yeow et al., 2025) or *in vitro* binding experiments (Fig. 1D). This indicates that EB1 is not a physiological substrate of TRIM37, despite their spatial proximity within the cell. We therefore used EB1 as a neutral scaffold to test whether appending the IDR+ASH8 domain would be sufficient to promote TRIM37-dependent degradation (Fig. 4A). Using a tetracycline-inducible system in human colorectal cells, we expressed GFP-EB1 or GFP-EB1-IDR+ASH8 (Fig. 4B-D, condition 1) and then withdrew tetracycline to monitor protein stability in the presence or absence of TRIM37, via siRNA-mediated knockdown (Fig. S3A). After tetracycline removal, GFP-EB1 levels increased slightly, whereas GFP-EB1-IDR+ASH8 levels decreased, indicating degradation (Fig. 4B-D condition 2). Depletion of TRIM37 prevented this decrease, restoring GFP-EB1-IDR+ASH8 levels to those of unmodified EB1 (Fig. 4B-D condition 3). TRIM37 depletion had no detectable effect on GFP-EB1 levels. Thus, TRIM37 mediates degradation of EB1 only when it is fused to the IDR+ASH8 module. We conclude that the IDR+ASH8 degron is sufficient to convert a non-targeted protein into a TRIM37 substrate.

**Figure 4.**
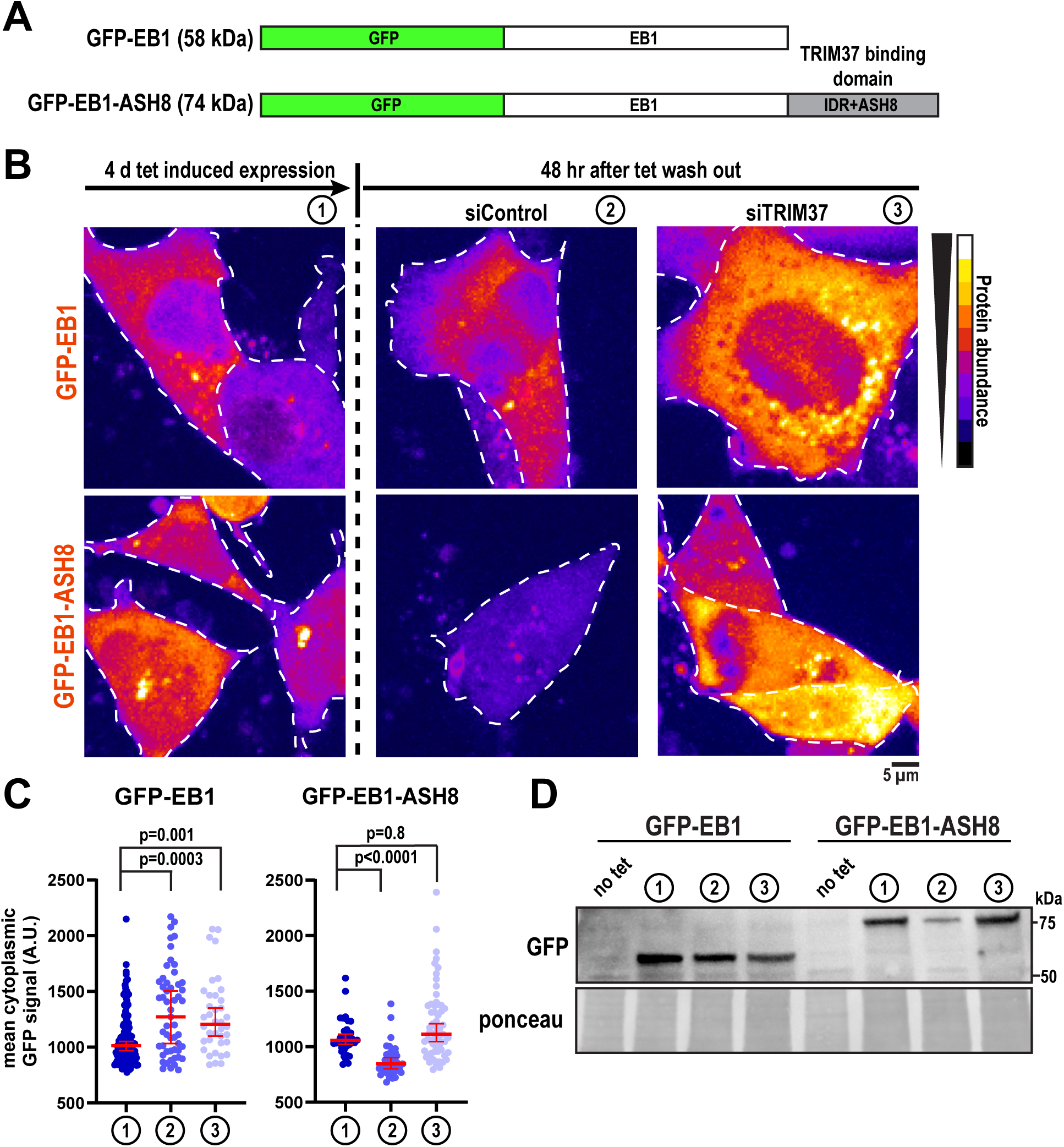
IDR+ASH8 functions as a transferable degron recognized by TRIM37. A) Diagram of EB1 constructs expressed in DLD-1 cells. B) Representative images showing EB1 constructs: (1) during tetracycline-induced expression, (2) after tetracycline withdrawal with siControl, and (3) after tetracycline withdrawal with siTRIM37. Cells are outlined in white. C) Quantification of cytoplasmic EB1-construct fluorescence under conditions described in (B) (median ± 95% CI; >35 cells, 2 biological replicates). P values from Kruskal-Wallis test, followed by Dunn’s multiple comparison test. D) Western blot against GFP showing total EB1-construct abundance under the conditions in (B).

### Biochemical analysis of the TRIM37-Cep192(IDR+ASH8) complex

We next sought to define the structural mechanism by which TRIM37 recognizes the IDR+ASH8 module. The only known substrate motif recognized by TRIM37 is a C-terminal disordered peptide motif within Centrobin (Bellaart et al., 2025). In contrast, AlphaFold3 predicts a tight beta barrel structure for ASH8 (Fig. 5A). Thus, we hypothesized that TRIM37 has multiple modes of substrate recognition.

**Figure 5.**
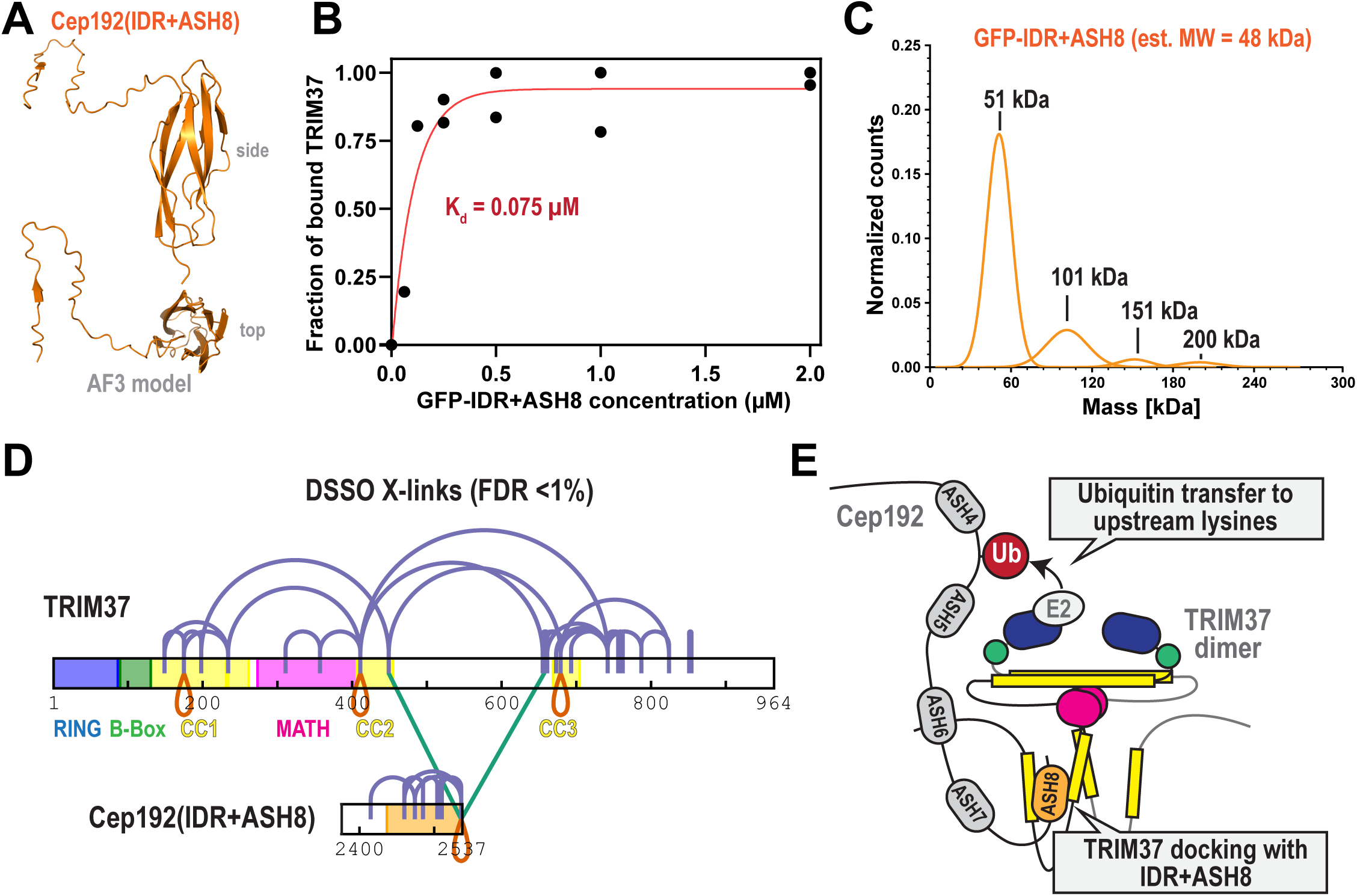
Structural analysis of the TRIM37-IDR+ASH8 complex. A) AlphaFold3 prediction of the IDR+ASH8 domain of Cep192. Side and top views shown. pTM = 0.68 for the whole domain; pTM = 0.86 for ASH8. B) Purified GFP-IDR+ASH8 was incubated with 30 nM TRIM37, then the complex was precipitated. The amount of TRIM37 remaining in the supernatant was measured to calculate the fraction of bound TRIM37 with each concentration of GFP-IDR+ASH8. K_d_ was determined from a one phase association fit (red line; R^2^ = 0.93). n = 2 biological replicates. Associated with Fig. S3B, C. C) Mass photometry of 20 nM GFP-IDR+ASH8. The data represent three biological replicates. D) DSSO crosslinking map of TRIM37 and IDR+ASH8. ASH8 is colored orange; intrinsically disordered regions are colored white. Cross-links: purple, intramolecular; green, intermolecular; red, intermolecular links between the same amino acid. n = 3 biological replicates, calculated FDR <1%. Associated with Fig. S3D. E) Model for how TRIM37 dimers recognize and ubiquitinate the C-terminus of Cep192.

We first measured binding affinity. We incubated 30 nM TRIM37 with increasing concentrations of GFP-IDR+ASH8 and GFP-Trap beads. After 1 hr, we spun down the beads and measured the concentration of unbound TRIM37 that remained in the supernatant (Fig. S3B, C) (Pollard, 2010). Plotting the bound fraction of TRIM37 vs. IDR+ASH8 concentration revealed binding occurred with mid-nanomolar affinity (K_d_ = 75 nM; Fig. 5B). This value is ∼180-fold lower than the K_d_ of TRIM37 binding with the Centrobin peptide motif (K_d_ = 1400 nM, (Bellaart et al., 2025)). Mass photometry revealed that GFP-IDR+ASH8 is largely monomeric but can form small populations of dimers, trimers, and tetramers (Fig. 5C). This result indicates that TRIM37 can recognize a non-oligomeric domain with high affinity.

Next, we performed cross-linking mass spectrometry (XL-MS) to map the interactions between TRIM37 and IDR+ASH8. We used 1 mM DSSO, an amine-reactive, MS-cleavable molecule that links lysine residues. MS analysis with data filtering (<1% calculated false discovery rate (FDR), n= 3 replicates) identified 74 total cross-links with high confidence: 55 specific to TRIM37 molecules, 19 specific to IDR+ASH8, and 2 between TRIM37 and IDR-ASH8 (Fig. 5D). When overlaid onto AlphaFold3 models, all cross-links within IDR+ASH8 satisfied distance constraints expected for DSSO (<35 Å; Fig. S3D), supporting that ASH8 conforms to a compact beta barrel structure. An intermolecular self-link (K2536-K2536; red loop in Fig. 5D) indicates that IDR+ASH8 can at least form dimers, consistent with mass photometry. We also identified three self-links within TRIM37 (K175-K175, K411-K411, K679-K679), consistent with TRIM37 forming dimers in cells (Fig. 5D) (Yeow et al., 2025). In Fig. S3D, we show cross-links that satisfy distance constraints when mapped onto an AlphaFold3 model of the dimeric N-terminus (a.a. 1-458), which has well defined structural motifs. These results support the accuracy of our cross-link assignments.

XL-MS revealed that the ASH8 beta barrel interacts with the coiled-coil domain (CC2) following the MATH domain and a predicted coiled-coil domain (CC3) in the tail of TRIM37 (Fig. 5D; FDR<1%). By choosing a medium confidence cutoff (FDR<5%), we revealed an additional interaction between CC3 and part of the IDR just upstream of ASH8 (Fig. S3E). These data are consistent with our biochemical results showing that both the IDR and ASH8 are needed for TRIM37 binding (Fig. 1). This differs from Centrobin, which was shown to bind the MATH domain of TRIM37 (Bellaart et al., 2025). Furthermore, we identified numerous high-confidence cross-links within CC3, suggesting that this region may represent a new structural motif. Cross-links between CC3, the intrinsically disordered tail, and CC2 indicate that TRIM37’s tail (a.a. 459-964) interacts with the N-terminus. Our data support a model whereby TRIM37 uses two separate coiled-coil motifs to bind the bipartite IDR+ASH8 motif with high affinity.

## DISCUSSION

How E3 ubiquitin ligases target specific proteins is an important but poorly understood area of cell biology. In this study, we revealed the mechanism by which the E3 ubiquitin ligase TRIM37 targets Cep192, a protein essential for centrosome assembly and function. We demonstrated that TRIM37 directly engages Cep192 through a defined C-terminal module called IDR+ASH8. TRIM37 then ubiquitinates an upstream cluster of 7 lysine residues on Cep192 (Fig. 5E). Ubiquitination reduces the stability of Cep192, which regulates Cep192 steady state levels and centrosome size. Finally, we showed that the IDR+ASH8 module can transform EB1 into a TRIM37 substrate, establishing that IDR+ASH8 is a degron. Our findings thus provide a structural mechanism for how TRIM37 contributes to centrosome homeostasis.

Our data demonstrate that TRIM37-mediated ubiquitination is required for Cep192 degradation, and that this process depends on both the TRIM37-binding region and the lysines that accept ubiquitin. In cells, eliminating the TRIM37-interaction site (ΔASH8) abolished degradation and completely prevented TRIM37 binding, showing that access to this motif is essential for initiating the reaction. Likewise, substituting the 7 ubiquitinated lysines with arginines (7KtoR) stabilized Cep192 and blocked its ubiquitination, establishing that these residues are required for substrate degradation. Notably, although 7KtoR still engaged TRIM37, its reduced binding compared with the WT protein suggests that lysine availability could regulate ligase association. This decreased affinity further supports the idea that ubiquitination weakens TRIM37–Cep192 interactions. Such ubiquitin-triggered substrate release is consistent with established mechanisms for other RING E3s, including SCF(Cdc4) and APC/C (Deffenbaugh et al., 2003; Sitry-Shevah et al., 2024).

Overexpression of both Cep192 mutants (7KtoR and ΔASH8), but not Cep192(WT), significantly prolonged mitosis. A previous study showed that mitotic duration is moderately extended in *TRIM37* knockout cells (Meitinger et al., 2021), while another study showed that mitotic duration is not affected (Tkach et al., 2022). Thus, it is difficult to conclude if the mitotic delay phenotype we observed results from TRIM37’s inability to ubiquitinate Cep192 or from another cause. Unexpectedly, both mutants exhibited reduced Cep192 fluorescence intensity at mitotic centrosomes, indicating that ubiquitination-deficient Cep192 forms a less robust PCM scaffold. In contrast, during interphase, both ΔASH8 and 7KtoR mutants accumulated more strongly at centrosomes than wild-type Cep192, suggesting that the need for ubiquitination is specific to mitotic PCM formation. Overexpression of PCM proteins might result in a multipolar spindle that extends the duration of mitosis. However, we observed that most examined cells expressing Cep192 constructs formed regular bipolar spindles (Fig. 3C, S2B). Thus, this mitotic phenotype is likely not caused by irregular spindle formation. Although unexpected, this finding raises the intriguing possibility that Cep192 ubiquitination acts as an important post-translational signal that supports correct PCM maturation during mitosis. This could impact cell proliferation given that mitotic PCM assembly affects the speed and fidelity of mitotic division in many species (Cabral et al., 2019; Conduit et al., 2014; Hamill et al., 2002; Lawo et al., 2012; Meitinger et al., 2020). These data align with earlier reports of increased levels of ubiquitinated proteins at the centrosome in metaphase relative to S phase (Kimura et al., 2014). Together, these observations point to regulatory layers that extend beyond the scope of the present study and warrant further investigation.

While the functional consequences of ubiquitination-deficient Cep192 are intriguing, the central focus of this work is to define how TRIM37 identifies its substrates. We found that TRIM37 recognizes a specific module comprising the ASH8 domain and its adjacent linker (IDR+ASH8). Moreover, we demonstrated that the fusion of the IDR+ASH8 module to an unrelated protein, EB1, is sufficient to trigger its degradation via TRIM37. AlphaFold3 predicts that the IDR region is not structured (Abramson et al., 2024) and that the ASH8 domain conforms to a compact beta barrel (35Å length, 12 Å width, and 39 Å in diameter). Interestingly, several TRIM37 interactors identified by proximity labeling (Yeow et al., 2020; Yeow et al., 2025) are predicted to contain similar structures: Cep170B, C2CD3, RPGRIP1L, TJP1 and MIS18BP1. This raises the possibility that a compact beta barrel could be a general targeting motif for TRIM37. However, simply having a compact beta barrel is not sufficient, since full length Cep192 contains 8 beta barrels (all ASH-like domains), but only the last one engages TRIM37. Structural comparisons of predicted ASH8 and ASH6 with the solved ASH7 structure (PDB: 6FVI) reveal that ASH8 lacks several protrusions found in the others (Fig. S3F). Moreover, ASH8 has a more positively charged surface (Fig. S3G). Identifying the functional epitope within ASH8 will be crucial for defining these structural distinctions that permit TRIM37 recognition. This pursuit would be aided by identifying additional bona fide substrates of TRIM37 and determining the similarities between their docking motifs.

Our work raises the compelling possibility that TRIM37 has multiple modes of substrate recognition tailored for specific functions. Recent work demonstrated that a small peptide motif mediates the interaction between TRIM37 and Centrobin to prevent the formation of pathological PCM-like condensates (Balestra et al., 2021; Bellaart et al., 2025; Meitinger et al., 2021). In this case, TRIM37 uses its MATH domain to weakly bind (K_d_ = 14 μM) Centrobin’s peptide motif. It was hypothesized that weak binding was a key feature of this regulatory system. Large-scale multimerization of the substrate would create a high local concentration of binding sites, thus attracting TRIM37 through an avidity effect. This would also promote TRIM37 multimerization, which is known to increase its ligase activity (Bellaart et al., 2025). By this mechanism, TRIM37 would only ubiquitinate and destroy aggregated forms of Centrobin. In contrast, our results demonstrate that TRIM37 can interact with a non-multimeric substrate: the IDR+ASH8 domain of Cep192. Our XL-MS results showed that two separate coiled-coil domains (CC2, CC3) on TRIM37 connect with IDR+ASH8 (Fig. 5D), suggesting that the MATH domain is not involved. We speculate that this mode of substrate recognition regulates steady state levels or activities of functional proteins. Centrobin contains no beta barrels, while the C-terminus of Cep192 does not contain the same IDR found in Centrobin. These data are consistent with a model whereby TRIM37 has two modes of substrate docking: 1) low-affinity, MATH domain-mediated recognition of a disordered peptide motif, and 2) high-affinity, coiled-coil-mediated recognition of an ASH8-like beta barrel with a preceding IDR. We speculate that TRIM37 may switch between these modes depending on cellular context, enabling it to regulate diverse substrates with different structural features and functional consequences.

Together, our findings provide a new mechanism for TRIM37 substrate recognition, revealing that the ligase can engage clients through high-affinity binding to a C-terminal bipartite motif. The IDR+ASH8 module of Cep192 acts as a transferable degron, explaining how TRIM37 regulates Cep192 levels and potentially other substrates. Given the high sequence conservation of Cep192 and TRIM37 amongst vertebrates, this mechanism of PCM protein turnover may be broadly conserved. Moreover, our discovery that the IDR+ASH8 module functions as a TRIM37-specific degron has broader implications for the hundreds of TRIM-family and other RING-type E3 ligases for which substrate recognition rules have yet to be defined (Zhang et al., 2025). The portability of this bipartite degron suggests that similar mixed-architecture motifs may exist across the proteome but remain unrecognized.

## FUNDING

J.B. Woodruff is supported by supported by a Cancer Prevention and Research Institute of Texas grant (RR170063), the Welch Foundation Grant (V-I-0004-20230731), an R35 grant from the National Institute of General Medical Sciences (5R35GM142522), and the Endowed Scholars program at UT Southwestern. Some data presented in this report were acquired with a mass photometer that was supported by award S10OD030312-01 from the National Institutes of Health.

## ACKNOWLEDGEMENTS

We thank the UT Southwestern Proteomics Core Facility for mass spectrometry and cross-linking analysis; Andrew Holland and Peter Ly for providing cell lines and reagents; Vincent Tagliabracci and Mike Henne for providing additional molecular biology reagents. We thank Manolo Rios for helping with cross-linking mass spectrometry.

## Author contributions

W.E. Stachera designed, performed, and analyzed the majority of the experiments. J. Tafur expressed and purified Cep192 protein fragments. N.E. Familiari expressed full-length Cep192 and TRIM37(C18R) proteins in insect cells using baculovirus. K. Yaguchi performed immunofluorescence experiments. J.B. Woodruff performed mass photometry and protein affinity assay and supervised the study. W.E. Stachera wrote the manuscript with input from J.B. Woodruff.

## DATA AVAILABILITY

Mass spectrometry raw data are accessible from MassIVE. Accession number: MSV000100205. Link for reviewers to download data: ftp://MSV000100205@massive-ftp.ucsd.edu

Password: UTSW2025

## MATERIALS AND METHODS

### Protein expression and purification

Cep192 constructs were inserted into a baculoviral expression plasmid (pOCC29) using standard restriction cloning. Baculoviruses were generated using the FlexiBAC system (Lemaitre et al., 2019) in SF9 cells. We used SF9-ESF *Spodoptera frugiperda* insect cells (Expression Systems) grown at 27°C in ESF 921 Insect Cell Culture Medium (Expression Systems) supplemented with Fetal Bovine Serum (2% final concentration). Protein was harvested 72 hr after infection during the P3 production phase. Cells were collected, washed, and resuspended in harvest buffer (25 mM HEPES, pH 7.4, 150 mM NaCl). For the expression of smaller Cep192 truncations and EB1 constructs, we used BL21(DE3) bacterial cells. All expression plasmids are listed in Table S1 and S2.

All subsequent protein purification steps were performed at 4°C. Cell pellets were resuspended in Lysis Buffer (25 mM HEPES, pH 7.4, 30 mM imidazole, 500 mM KCl, 0.5 mM DTT, 1% glycerol, 0.1% CHAPS) + protease inhibitors and then lysed using a dounce homogenizer. Proteins were bound to Ni-NTA (Qiagen), washed with 10 column volumes of Lysis Buffer, and eluted with 250 mM imidazole. The eluates were concentrated using Ultra-15 Centrifugal Filters (Millipore) and incubated with 3C PreScission/ TEV protease overnight to remove purification tags. Then we ran purified proteins through AKTA FPLC using Superdex^TM^ 200 (Cytiva, 28-9909-44), or Superdex^TM^ 75 (Cytiva, 29148721) Increas 10/300 GL column, depending on the size of protein. During gel filtration all proteins were in Gel Filtration Buffer (25 mM HEPES, pH 7.4, 500 mM NaCl, 0.5 mM DTT, 1% glycerol, 0.1% CHAPS). All eluted proteins were then concentrated and aliquoted in PCR tubes, flash-frozen in liquid nitrogen, and stored at −80°C. Protein concentration was determined by measuring absorbance at 280 nm using a NanoDrop ND-1000 spectrophotometer (Thermo Fisher Scientific).

### Ubiquitination assay and detection by mass spectrometry

We combined ligases previously associated with TRIM37 (Bhatnagar et al, 2014): 10 nM E1 UBE1 (Biotechne, E-305-025) and 10 nM E2 UbcH5b/UBE2D2 (Biotechne, E2-622-100). We added 500 nM HA-Ubiquitin (gifted from the Tagliabracci lab), purified Cep192 (1 µM) protein and ubiquitin reaction buffer (5 mM MgCl_2_, 0.5 mM DTT, 2mM ATP, 25 mM HEPES, 100 mM KCl, 100nM ZnCl_2_). To the positive sample we added 1 µg Trim37 (Abcam ab269066), while the negative sample did not contain any E3 ligases. We sent the samples for identification of post translational modification, specifically for GG marks that are left after ubiquitination.

Samples were digested overnight with trypsin (Pierce) following reduction and alkylation with DTT and iodoacetamide (Sigma–Aldrich). Following solid-phase extraction cleanup with an Oasis HLB µelution plate (Waters), the resulting peptides were reconstituted in 10 uL of 2% (v/v) acetonitrile (ACN) and 0.1% trifluoroacetic acid in water. 5 uL of this was injected onto an Orbitrap Eclipse mass spectrometer (Thermo) coupled to an Ultimate 3000 RSLC-Nano liquid chromatography system (Thermo). Samples were injected onto a 75 μm i.d., 75-cm long EasySpray column (Thermo), and eluted with a gradient from 0-28% buffer B over 90 min. Buffer A contained 2% (v/v) ACN and 0.1% formic acid in water, and buffer B contained 80% (v/v) ACN, 10% (v/v) trifluoroethanol, and 0.1% formic acid in water. The mass spectrometer operated in positive ion mode with a source voltage of 2.5 kV and an ion transfer tube temperature of 300 °C. MS scans were acquired at 120,000 resolution in the Orbitrap and up to 10 MS/MS spectra were obtained in the Orbitrap for each full spectrum acquired using higher-energy collisional dissociation (HCD) for ions with charges 2-7. Dynamic exclusion was set for 25 s after an ion was selected for fragmentation. Each sample was injected twice.

Raw MS data files were analyzed using Proteome Discoverer v.3.0 (Thermo), with peptide identification performed using Sequest HT searching against the human reviewed protein database from UniProt (downloaded January 4, 2024, 20354 entries). Fragment and precursor tolerances of 10 ppm and 0.6 Da were specified, and three missed cleavages were allowed. Carbamidomethylation of Cys was set as a fixed modification, with oxidation of Met, and GlyGly modification of Lys set as variable modifications. The false-discovery rate (FDR) cutoff was 1% for all peptides. The sum of peak intensities for all peptides matched to a protein were used as protein abundance values. A list of identified peptides together with PTM sites found can be found in Extended Data Set 1.

### Cross-linking mass spectrometry (XL-MS)

Reactions were assembled using purified proteins at final concentrations of 5 µM TRIM37(C18R) (Table S1) and 5 µM IDR+ASH8 (Table S2). The proteins were combined with 100 nM ZnCl₂, 25 mM HEPES, and 150 mM KCl (pH 7.4). Reactions were incubated for 20 min at 37 °C with shaking at 350 rpm. Cross-linking was initiated by adding DSSO to a final concentration of 1 mM, followed by a 5-min incubation. The reaction was then quenched with ammonium bicarbonate (ABC; 50 mM final) and incubated for an additional 30 min at 37 °C with shaking at 350 rpm. Samples were denatured by adding 6× SDS sample buffer and heating for 5 min at 95 °C. Proteins were separated by SDS– PAGE, stained with Coomassie Blue, and the band corresponding to the TRIM37–ASH8 complex was excised. Gel slices were submitted for mass spectrometry analysis, requesting three injections per sample.

Samples were digested overnight with trypsin (Pierce) following reduction and alkylation with DTT and iodoacetamide (Sigma–Aldrich). Following solid-phase extraction cleanup with an Oasis HLB µelution plate (Waters), the resulting peptides were reconstituted in 10 µL of 2% (v/v) acetonitrile (ACN) and 0.1% trifluoroacetic acid in water. 3 aliquots of each sample were separately injected onto an Orbitrap Fusion Lumos mass spectrometer coupled to an Ultimate 3000 RSLC-Nano liquid chromatography system. Samples were injected onto a 75 um i.d., 75-cm long EasySpray column (Thermo) and eluted with a gradient at a flow rate of 250 nL/min from 0-5% buffer B over 1 min, 5%-40% B over 60 minutes, 40%-99% over 25 minutes, and held at 99% B for 5 minutes before returning to 0% B for column equilibration. Buffer A contained 2% (v/v) ACN and 0.1% formic acid in water, and buffer B contained 80% (v/v) ACN, 10% (v/v) trifluoroethanol, and 0.1% formic acid in water. The mass spectrometer operated in positive ion mode with a source voltage of 2.0 kV and an ion transfer tube temperature of 275 °C. MS scans were acquired at 120,000 resolution in the Orbitrap. MS/MS spectra were obtained in the Orbitrap using top-speed mode (5 sec) for each full spectrum acquired using collisionally-induced dissociation (CID) for ions with charges 3-7. Dynamic exclusion was set for 25 s after an ion was selected for fragmentation. MS2 fragment ions with 1-4 charge states and mass difference of 31.9721 with 10-100% intensity range were selected for MS3 CID fragmentation, with a mass tolerance of 25 ppm, 2 m/z isolation window, and at 15,000 resolution.

Raw MS data files were analyzed using Proteome Discoverer v.3.0 (Thermo), with peptide identification performed using MS Annika 2.0 (Birklbauer et al., 2023) searching against the amino acid sequences of TRIM37 and CEP192. Fragment and precursor tolerances of 10 ppm (MS1), 20 ppm (MS2), and 20 ppm (MS3) were specified, and up to four missed cleavages were allowed. DSSO (+158.004 Da, modification on K) was set as the crosslinker modification, with N-terminal crosslinks allowed. Diagnostic ions of 138.0911, 155.1179, 170.0634, and 187.0900 were included. Carbamidomethylation of Cys was set as a fixed modification, with oxidation of Met and DMSO modification of Lys set as a variable modification. The false-discovery rate cutoff was 1% (i.e., estimated FDR) at the CSM and cross-link levels. The peak intensities of the peptides were used as abundance values. Calculated FDRs ranged from 5-7% for all cross-links (see Extended Data Set 2). In Figure 5, the most confident cross-links (calculated FDR <1%) were chosen and visualized with xiView (https://www.xiview.org/index.php).

### Western Blotting

Samples were incubated with SDS sample buffer for 5 min at 95°C and separated by SDS-PAGE. Protein from each gel was transferred to a nitrocellulose membrane using a Trans-blot turbo transfer for high molecular weight proteins (10 min). Membranes were incubated in a blocking buffer consisting of 1× TBS-T + 3% Blotting-Grade Blocker (BioRad) shaking at room temperature for 1 hr. Membranes were then washed three times with fresh 1× TBS-T and incubated shaking with primary antibodies overnight at 4°C. The primary antibody was washed three times with fresh 1× TBS-T and incubated by shaking with secondary antibodies at room temperature for 1 hr. Each membrane was then incubated in ECL reagent (Thermo Fisher Scientific SuperSignal West Femto) for 5 min and imaged with a ChemiDoc Touch Imaging System. Primary antibodies: Mouse anti-alpha tubulin (Cell Signaling Technologies, 3873S, 1:1,000); Goat anti-enhanced GFP (1:5,000; Dresden PEP facility, 15 mg/ml; Poser et al., 2008), Rabbit anti-TRIM37 (1:1000; Bethyl/ Thermofisher A301-174A); Secondary Antibodies (1:20,000-1:100,000): HRP conjugated Goat anti-Rabbit IgG (1 mg/ml) (Invitrogen, 65-6120); HRP conjugated Goat anti-Mouse (1.5 mg/ml) (Invitrogen, 62-6520); and HRP conjugated Donkey anti-Goat (1 mg/ml) (Invitrogen, A15999).

### Co-immunoprecipitation

Purified Cep192 constructs (1µM) and EB1 (1µM, control) were incubated with TRIM37 (1 µg) for 1hr at 37°C with rotation. GFP-Trap® Magnetic Particles M-270 (chromotek, gtd-20) or Dynabeads ™ Protein A (conjugated with TRIM37 antibody, Bethyl A301-174A) were pre-washed with buffer (150 mM KCl, 25 mM HEPES, 1% Glycerol, 0.05% NP-40). The TRIM37-GFP protein reactions were incubated with 25 µl prewashed beads at room temperature for 10 min with rotation, followed by incubation at 4°C for 1 hr with rotation. Then beads were washed 3 times with washing buffer followed by one additional wash with a higher salt washing buffer (500 mM KCl, 25 mM HEPES, 1% Glycerol, 0.05% NP-40). We eluted the bound fraction by incubating with SDS sample buffer at 95°C for 5 min and the bound components were detected by western blotting.

### Cell culture

DLD-1 cells (CCL-221) were cultured in DMEM (Corning 10-017-CV) supplemented with 10% fetal bovine serum (Gibco 10082147) and 100 U/mL penicillin-streptomycin (Sigma P4333). All cell lines were maintained at 37°C in a humidified incubator with 5% CO₂ and 21% oxygen. The DLD-1 cell line with the Flp-In™ T-REx™ system was generously provided by the Peter Ly laboratory (Tighe et al., 2004).

### Cell lysis

For each pull-down experiment, cells were harvested from 10-cm plates, counted, and 8 × 10⁶ cells were used per condition. Cells were washed once with ice-cold PBS supplemented with protease inhibitor cocktail (Millipore, 539134-1SET). Cell lysis was performed using a lysis buffer composed of 500 mM KCl, 25 mM HEPES (pH 7.4), 1% glycerol, and 1% NP-40. We sonicated cells for 15 seconds using 30% amplitude.

### Generation of cell lines expressing protein constructs

Cep192 (cDNA clone NM_032142.3; GenScript) and EB1 (Woodruff et al., 2017; Zanic et al., 2009) constructs were made via PCR amplification verified through Sanger sequencing. All plasmids used to transfect DLD-1 cell line are listed in Table S3. Stable, isogenic cell lines overexpressing Cep192 or EB1 variants were established by co-transfecting the pcDNA5 plasmids with the pOG44 FLP recombinase vector using JetPrime™ reagent. Positive clones were selected with 100 µg/mL hygromycin (Millipore Sigma), and construct expression was induced by treating cells with 2 µg/mL tetracycline (Millipore Sigma). For monitoring construct stability, we first overexpressed each construct for 96 hr by adding tetracycline to the media (2 µg/mL). After imaging the 96 hr expression time point, we washed out cells 3 times with PBS to remove tetracycline completely. We imaged cells again 24 hr, and 48 hr after tetracycline wash out to track the construct degradation process.

### RNA interference

For TRIM37 silencing, DLD-1 cells were transfected with siTRIM37 (Thermofisher, siRNA ID 144318) or Stealth™ Negative Control (Invitrogen, 452002) using Lipofectamine™ RNAiMAX reagent.

### Imaging

Cells were seeded onto a 24-well glass-bottom tissue culture plate (Celltreat, 229125) and then grew for 48 hr before imaging. Each cell line was imaged in DMEM without phenol red (Gibco 21063-029), and DNA was stained overnight with 100 nM SiR-DNA dye (Cytoskeleton CY-SC007) before imaging. Live-cell imaging was performed inside a stage-top incubator (Tokai Hit STX), maintained at 37°C with 5% CO₂. Images were acquired using an inverted Nikon Eclipse Ti2-E microscope equipped with a Yokogawa CSU-W1 confocal scanner unit, piezo Z stage, and an iXon Ultra 888 EMCCD camera (Andor), controlled by Nikon Elements software.

A 60× 1.2 NA Plan Apochromat water-immersion objective was used to acquire 31 Z-stacks (0.5-µm steps) each with 100 ms laser exposure. Fluorescence imaging was performed sequentially using the following settings:

- 488 nm excitation (15% laser power, 2 × 2 binning)
- 561 nm excitation (15% laser power, 1 × 1 binning)
- 640 nm excitation (10% laser power, 1 × 1 binning)

For imaging mitotic progression, we first expressed constructs inside cells using tetracycline treatment for 24-72 hr. Then we arrested cells in late G2 phase by incubating them overnight with 9 µM CDK1 inhibitor (RO-3306, S7747). After washing out the drug, we waited an hour to make sure most cells would enter into mitosis. Then we imaged cells using 5% laser power with 1×1 binning.

### Image quantification and statistical analysis

Images were analyzed in FIJI using semi-automated, threshold-based particle analysis with the set area for every Cep192 foci or EB1 cytoplasmic signal. Data were plotted and statistical tests were performed using GraphPad prism. The sample size, measurement type, error type, and statistical test are described in the figure legends, where appropriate.

### Immunofluorescence

A day before tetracycline induction, cells were plated onto collagen-coated 24-well glass plates (Corning, CLS354236). To induce Cep192 overexpression, tetracycline was added to a final concentration of 2 µg/mL and cells were incubated for 96 hr. Cells were then washed once with PBS and fixed in 100% methanol for 15 min at –30 °C. After fixation, methanol was removed by rinsing with PBS and incubating the plate overnight at 4 °C in the dark. The following day, cells were permeabilized with 0.5% Triton X-100 for 5 min at room temperature and then blocked with 5% BSA in TBS for 30 min. Cells were incubated with primary antibodies conjugated with fluorophores for 48 hr at 4 °C in the dark: anti-Centrin-1 (Proteintech, CL594-12794; 1:500), and anti-GFP (ThermoFisher, A-21311; 1:500). After incubation, cells were washed twice with PBS, with a 10-min room temperature incubation during the second wash. Nuclei were stained overnight at 4 °C in the dark using DAPI (ThermoFisher, 62248; 1:1000 in PBS). Prior to imaging, cells were rinsed once with PBS.

### Cell viability assay

For each construct, cells were plated in triplicate on a 6-well plate at a density of 1 × 10⁵ cells per well. Tetracycline was added to the culture medium at a final concentration of 2 µg/mL to induce construct expression. We counted the number of cells on days 1, 3, and 6 using a hemocytometer, with all cells passaged back into culture after counting. The experiment was performed three independent times, and the mean cell number for each day of construct expression was plotted.

### qPCR

We used The Arcturus™ PicoPure™ RNA Isolation kit (Thermofisher, 12204-01) to purify RNA from the lysed cells expressing each construct for 96 hr, collected in triplicates. We measured RNA concentration using Nanodrop using 1 µg of RNA for further analysis. We performed cDNA synthesis from RNA using iScript Reverse Transcript Supermix for RT-qPCR (BioRad, 1708841) following the product protocol. Then we set up the RT-qPCR reaction using SsoAdvanced Universal SYBR® Green Supermix (BioRad, 1725271) and primers detecting GFP (Fw-GGACGACGGCAACTACAAGA, Rv – AAGTCGATGCCCTTCAGCTC) and housekeeping gene 36B4 as a control (Fw – AACATGCTCAACATCTCCCC, Rv-CCGACTCCTCCGACTCTTC) and following the product protocol.

### Mass photometry

GFP-Cep192(IDR+ASH8) was centrifuged to remove potential aggregates, then diluted into PBS to a final concentration of 20-50 nM on a clean glass coverslip. Protein was analyzed using a TwoMP mass photometer (Refeyn) using four species of BSA (monomer, dimer, trimer, and tetramer) to create a molecular weight standard curve.

### Protein affinity assay

Purified TRIM37 and GFP-Cep192(IDR+ASH8) were dialyzed and diluted using binding buffer (25 mM HEPES, pH 7.4, 150 mM KCl, 1 mM DTT). 30 nM TRIM37 was incubated with a range of GFP-Cep192(IDR+ASH8) concentrations, GFP-Trap® Magnetic Particles (Proteintech), and binding buffer to a final volume of 50 µl. After 1 hr incubation at room temperature, beads were sedimented, and the supernatant was collected and analyzed by SDS-PAGE and western blotting against TRIM37. Signal was measured using FIJI, the background signal subtracted, and remaining signal normalized to the negative control (no Cep192). The fraction of bound TRIM37 is defined as 1- (normalized supernatant signal).

---

**TABLE S1.** Baculovirus constructs protein expression.

| protein | sequence | plasmid name | parent plasmid | N-term tag | C-term tag |
| --- | --- | --- | --- | --- | --- |
| Cep192 | Full-length | JWV99 | pOCC29 | MBP-PreScission -<br>**eGFP** | PreScission-6xHis |
| Cep192 C-term | a.a. 2070-2537 | JWV106 | pOCC27 | MBP-PreScission | **eGFP**-PreScission-6xHis |
| TRIM37 C18R | Full-length | JWV130 | pOCC27 | MBP-PreScission | **eGFP**-PreScission-6xHis |

**TABLE S2.**
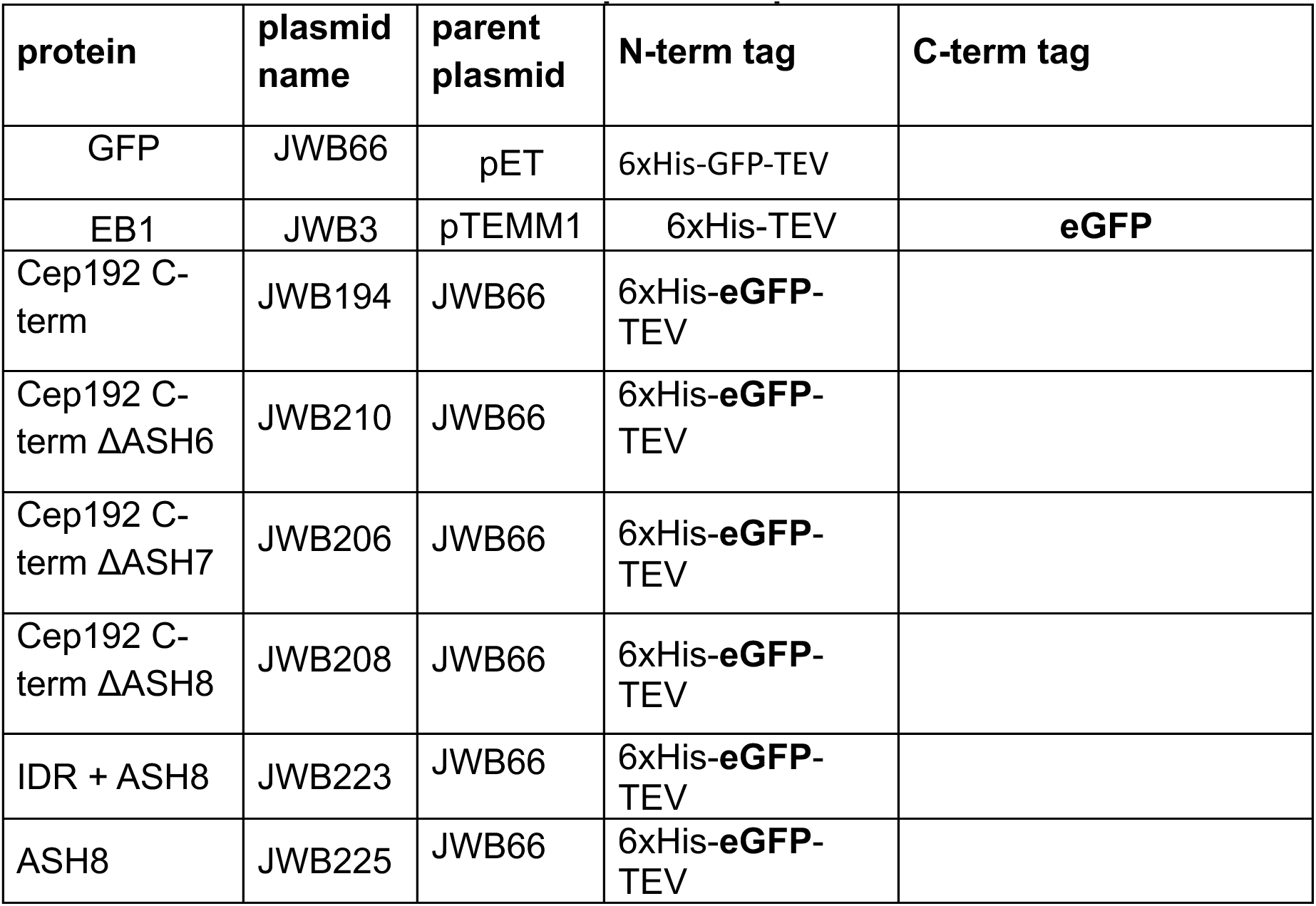
Bacterial constructs for protein expression.

**TABLE S3.** Constructs for protein expression using TREx Flp-In™ system in DLD-1 human colon cancer cells.

| Protein expressed | mutation | Cell line name | parent plasmid |
| --- | --- | --- | --- |
| GFP::Cep192 | WT | JWM25 | JWB179 |
| GFP::Cep192 | 7KtoR | JWM24 | JWB180 |
| GFP::Cep192 | no ASH8 (1-2437aa) | JWM28 | JWB224 |
| GFP::EB1 | WT | JMM 29 | JWB237 |
| GFP::EB1::ASH8 | IDR_ASH8 | JMM 30 | JWB230 |

**Extended Data Set 1. Mass spectrometry data for detection of ubiquitinated sites on Cep192.**

**Extended Data Set 2. Cross-linking mass spectrometry data.**

**Figure S1.**
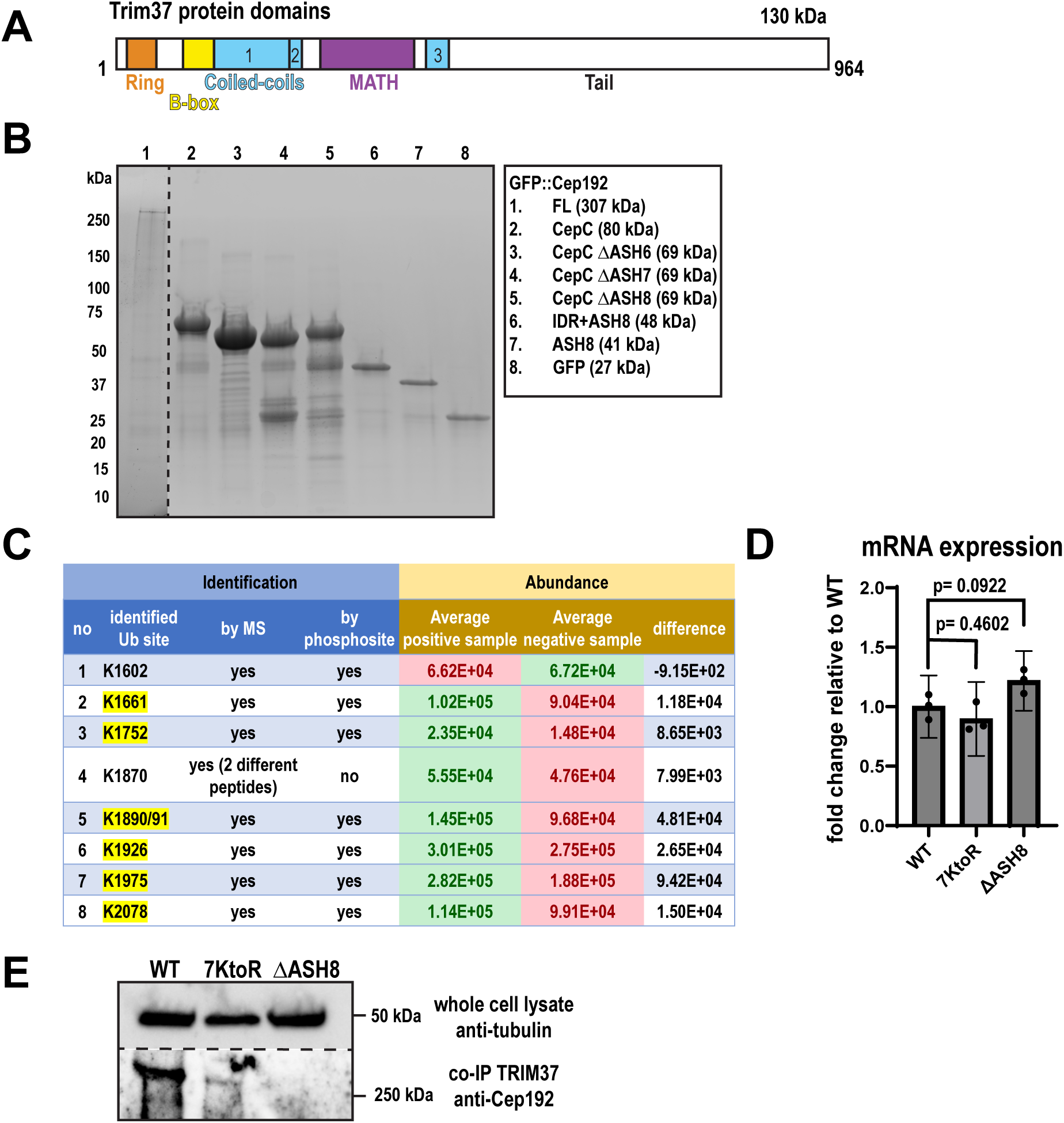
Identification of ubiquitinated residues and the TRIM37-binding site on Cep192. A) Domain architecture of TRIM37. B) SDS-PAGE gel showing purified full-length Cep192 and fragments. C) List of ubiquitinated lysines identified by MS and reported in PhosphoSitePlus (https://www.phosphosite.org/). Lysines highlighted in yellow were found to be ubiquitinated in our samples and in PhosphoSitePlus. D) qPCR analysis of construct expression across three biological replicates. Bars show mean with 95% CI; P values from one-way ANOVA followed Dunnett’s multiple comparison. E) Western blot showing binding efficiency of Cep192 constructs to TRIM37. Constructs were expressed inside DLD-1 cells and then pulled down using beads coated with TRIM37.

**Figure S2.**
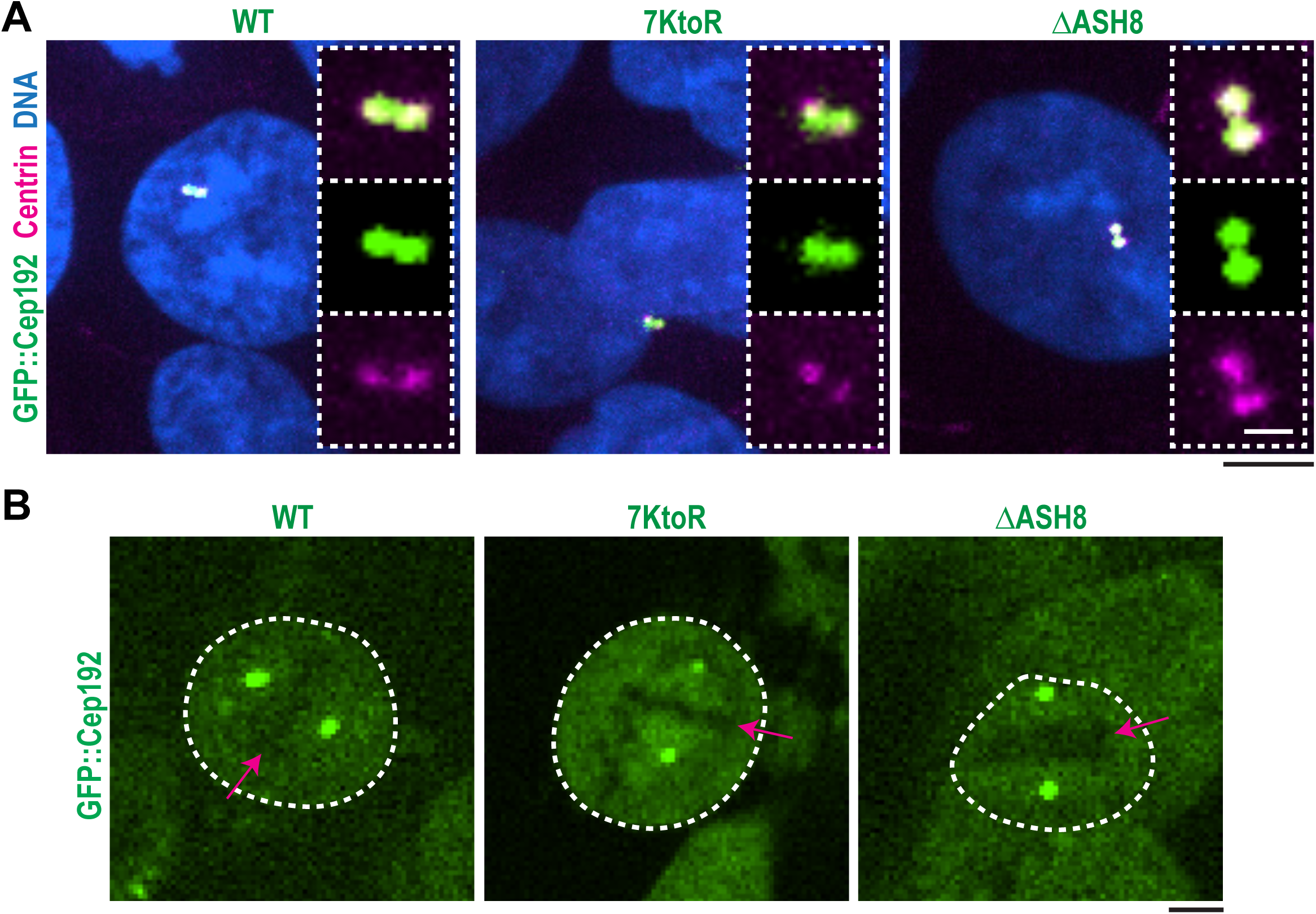
All Cep192 constructs localize to centrioles and support bipolar spindle formation. A) Immunofluorescence of cells after 96 hr of tetracycline-induced expression of Cep192 constructs. GFP constructs (green), Centrin (representing centrioles; magenta) and DNA (blue). Black scale bar: 5 µm; white scale bar: 1 µm. B) Representative images showing localization of GFP-Cep192 constructs in mitotic cells (outlined in white). Red arrows label chromosomes aligned at the metaphase plate. Scale bar: 5 µm.

**Figure S3.**
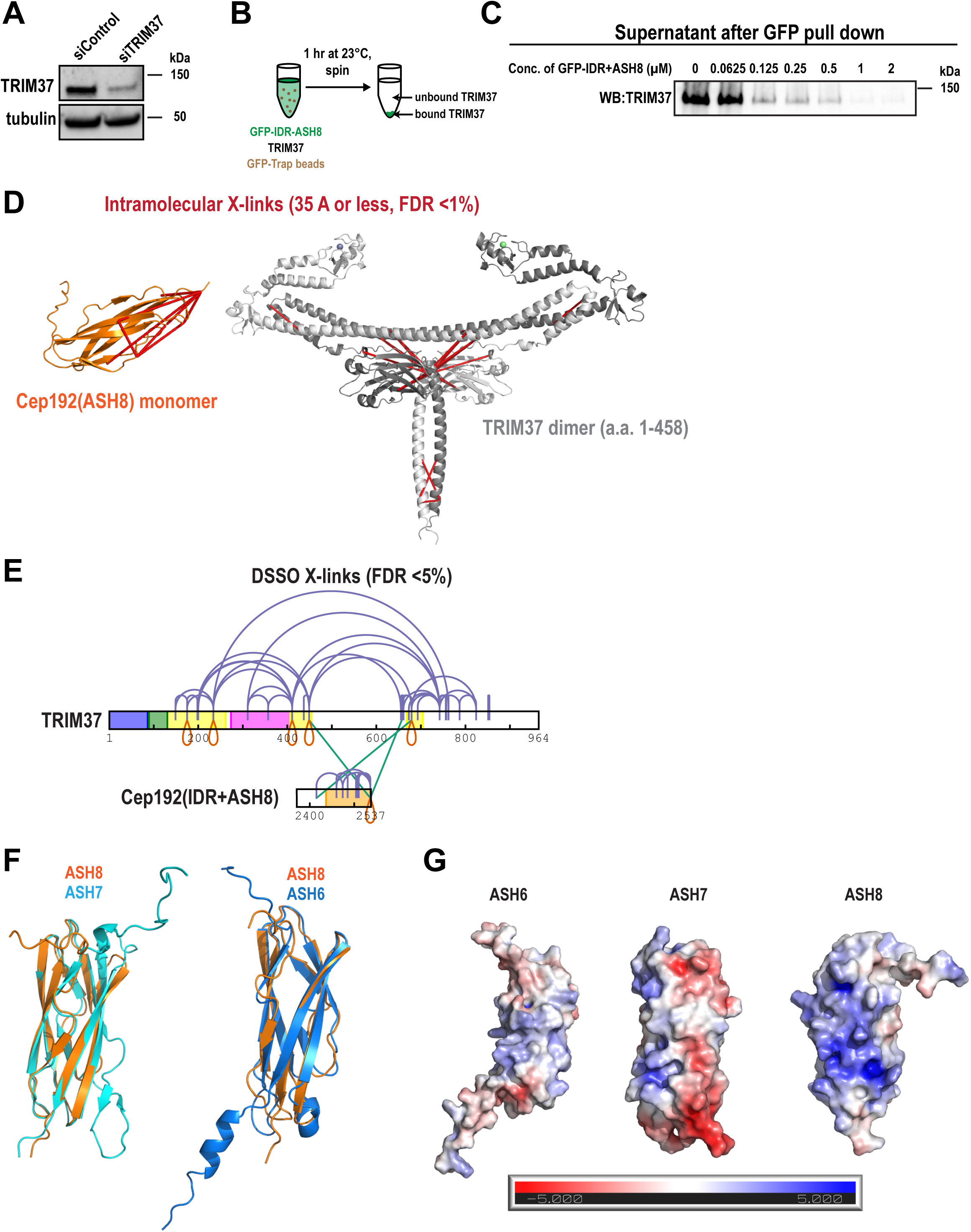
Biochemical characterization of the IDR+ASH8–TRIM37 interaction. A) Western blot against TRIM37 showing its abundance after siTRIM37 treatment. B) Schematic of pull-down assay from Fig. 5B. C) Western blot of TRIM37 found in the supernatant of experiments from Fig. 5B. D) Cross-links from Fig. 5D mapped onto AlphaFold3 predicted structures of the ASH8 domain and a TRIM37 dimer (a.a. 1-458). Shown are cross-links that satisfy the maximum distance constraint for DSSO (35 Å; between alpha carbons). E) DSSO crosslinking map of TRIM37 and IDR+ASH8 with calculated FDR <5% (medium confidence). ASH8 is colored orange; intrinsically disordered regions are colored white. Cross-links: purple, intramolecular; green, intermolecular; red, intermolecular links between the same amino acid. n = 3 biological replicates. F) Overlay of ASH8 (AlphaFold3 model) with ASH7 (PDB: 6FVI) and ASH6 (AlphaFold3 model). G) Estimated surface charge of structures in panel F (blue, positive; red, negative).

